# The vitellogenesis pathway of the tick host *Ixodes scapularis* regulates *Rickettsia buchneri* transcriptional state and persistence

**DOI:** 10.64898/2026.08.06.743197

**Authors:** Dattatray V. Sawant, Yating Dong, Haritha Katasani, Jonathan D. Oliver, Benjamin Cull, Benedict S. Khoo, Michel Shamoon-Pour, Amanda Baliban, Jianmin Zhong, Saravanan Thangamani, Ulrike G. Munderloh, Timothy J Kurtti, Xin-Ru Wang

**Affiliations:** Department of Microbiology and Immunology, Upstate Medical University, Syracuse, NY, 13210, USA; School of Life Sciences, Zhejiang Chinese Medical University, Hangzhou, China; School of Public Health, Division of Environmental Health Sciences, University of Minnesota, Minneapolis, MN, USA; Department of Entomology, University of Minnesota, Twin Cities, St. Paul, MN, 55108, USA; First-year Research Immersion, Binghamton University, Binghamton, NY, USA; Department of Anthropology, Binghamton University, Binghamton, NY, USA; Department of Biological Sciences, Humboldt State University, Arcata, California, USA; SUNY Center for Vector-Borne Diseases, Upstate Medical University, Syracuse, NY, 13210, USA; Institute for Global Health and Translational Sciences, Upstate Medical University, Syracuse, NY, 13210, USA

**Keywords:** *Ixodes scapularis*, tick endosymbiont, transovarial transmission, vitellogenesis, transcriptional quiescence, host-symbiont co-evolution, neofunctionalization, endosymbiont regulation, *Rickettsia buchneri*

## Abstract

Obligate endosymbionts relying on transovarial transmission must coordinate with host reproduction, yet the regulatory mechanisms remain poorly understood. Here we show that the vitellogenesis pathway of the tick *Ixodes scapularis* regulates the transcriptional state of its endosymbiont *Rickettsia buchneri* (*Rb*), separately from its abundance. In males, *Rb* DNA remained detectable, but bacterial transcription was strongly reduced across all examined genes, with markedly lower RNA/DNA ratios. In females, *Rb* was restricted to ovarian tissues (developing oocytes and interstitial cells), with no detection in salivary glands or midgut by TEM, FISH, or PCR. After blood feeding, *Rb* density within size-matched early-stage oocytes was significantly reduced. We further characterized two gene families mediating vitellogenesis: vitellogenin synthesis genes (Vgs, n = 20) and vitellogenin receptor genes (Vgr, n = 15). Vgs proteins showed conserved domain organization, whereas Vgr paralogs showed greater structural diversification. RNAi silencing of *Vgs20* or *Vgr12* altered *Rb* transcriptional profiles in both tick cells and ticks, although changes in bacterial load were not consistent between the two systems. Antibiotic depletion of *Rb* increased expression of both host genes. Together, these findings show that tick vitellogenesis pathways contribute to *Rb* regulation during reproduction and identify the *I. scapularis*-*Rb* system as a useful model for studying host control of obligate endosymbionts.

**IMPORTANCE:** Maternally inherited bacterial symbionts are commonly characterized by bacterial abundance. This study shows that bacterial abundance alone does not necessarily reflect symbiont functional state. In the blacklegged tick, the inherited symbiont *Rickettsia buchneri* persists in males but exhibits little transcriptional activity. In females, the host reproductive pathway determines whether the symbiont is active: silencing one component of this pathway changed bacterial gene expression without changing bacterial abundance. Hosts therefore regulate not only the abundance of inherited symbionts, but also their functional state. The same bacterial abundance can correspond to very different functional states. Understanding inherited symbioses therefore requires considering bacterial function alongside bacterial abundance.

## INTRODUCTION

Obligate endosymbionts that rely on vertical transmission must maintain stable associations with host reproductive systems for persistence across generations. This dependence links symbiont persistence directly to host reproduction. *Wolbachia*, found in an estimated 40-65% of insect species, has evolved multiple strategies to maximize maternal transmission (including cytoplasmic incompatibility, parthenogenesis induction, and male killing) to hijack host reproductive machinery for its own propagation (1–2). *Buchnera aphidicola* in aphids is compartmentalized within specialized bacteriocytes, where host-controlled mechanisms regulate symbiont density in response to developmental stage and nutritional demands (3–5). These examples demonstrate the close association between vertically transmitted endosymbionts and host reproductive biology. In hematophagous arthropods, vertically transmitted endosymbionts are widespread and ecologically important. However, the mechanisms of host regulation across the reproductive cycle remain poorly characterized, particularly in ticks where transovarial transmission differs from dipteran vectors.

*Ixodes scapularis*, the blacklegged tick, transmits multiple human pathogens in North America, including *Borrelia burgdorferi* (Lyme disease), *Anaplasma phagocytophilum* (anaplasmosis), and *Babesia microti* (babesiosis) (6). In contrast to other *Ixodes* species in Europe and Asia, where pathogenic spotted fever group rickettsiae are commonly detected, *I. scapularis* harbors no known pathogenic *Rickettsia*; instead, *Rickettsia buchneri* (*Rb*) is the only *Rickettsia* species in this tick (7). *Rb* is the dominant rickettsial symbiont in adult female *I. scapularis* and is maintained at high prevalence across natural populations through transovarial transmission (8–11). *Rb* encodes biosynthetic pathways for biotin and folate, nutrients absent from vertebrate blood, suggesting a potential nutritional contribution to the host (12). *Rb* has been shown to suppress the intracellular growth of pathogenic rickettsiae in tick cell culture (13), indicating that resident symbionts may influence pathogen establishment in ticks. The unique status of *Rb* as the sole *Rickettsia* species in this medically important vector, combined with its non-pathogenic nature and dependence on host reproduction, makes it a useful model for studying endosymbiont regulation in hematophagous arthropods. Yet the host mechanisms that regulate *Rb* activity and population dynamics across the reproductive cycle remain poorly understood. While our previous work demonstrated *Rb* association with autophagic compartments in ovarian interstitial cells, suggesting baseline autophagic regulation of the symbiont by the host (14), how this regulation extends across the full reproductive cycle, particularly during the physiological transition triggered by blood feeding, remains unknown.

In ticks, blood feeding triggers vitellogenesis and extensive ovarian remodeling. Vitellogenins are synthesized in peripheral tissues, secreted into the hemolymph, then internalized into developing oocytes via receptor-mediated endocytosis after blood feeding (15–16). These processes reshape the intracellular environment of developing oocytes. *Rb* has been observed to accumulate predominantly within developing oocytes (14), the primary cellular target of vitellogenin uptake, suggesting that vitellogenesis and symbiont maintenance may be coupled within the same cellular compartment. This possibility is supported by observations in other arthropod systems: *Wolbachia* in *Drosophila* exploits host vesicular trafficking and cytoskeletal networks to colonize oocytes (17), showing that host reproductive pathways can influence endosymbiont localization within oocytes. Whether host reproductive pathways regulate endosymbiont dynamics, rather than simply being co-opted by the symbiont, remains poorly characterized in tick systems. The vitellogenesis pathway therefore represents a direct and previously unexplored entry point to study endosymbiont regulation in *I. scapularis*.

Here, we address this question in *I. scapularis* through transcriptional, ultrastructural, genomic, and *in vivo* functional analyses. We find that the vitellogenesis pathway shapes *Rb* transcriptional state, a previously unrecognized regulatory role in arthropods. Moreover, bacterial load did not correlate with transcriptional state, showing that host control goes beyond simple population regulation. Our results establish the *I. scapularis-Rb* system as a model to study endosymbiont regulation in hematophagous arthropod vectors and suggest that reproductive pathways may play a broader role in symbiont control across vector systems. This regulation could be targeted to reduce tick reproductive fitness or vector competence, offering new approaches for tick-borne disease control.

## RESULTS

### *Rb* is transcriptionally silent in males

*Rb* was detected by PCR across all developmental stages of *I. scapularis*, including eggs, larvae, nymphs, and adults, using primers targeting the *gltA* gene, widely used for rickettsial detection, and the *ltrA2* gene, carried on an *Rb*-specific plasmid and absent from all other known *Rickettsia* species (Fig. 1a). At the individual level (n = 15 per group), *ltrA2* was detected in 33.3% (5/15) of unmated males and 66.7% (10/15) of mated males, compared to 100% (15/15) of females (Fig. 1b). Females showed significantly higher detection rates than both male groups (Fisher’s exact test, p < 0.05). We next quantified expression of seven *Rb* genes by RT-qPCR: *gltA*, *ltrA2*, *dnaK* (chaperone), *rpoB* (RNA polymerase β subunit), *groEL* (protein folding), *sca1* (surface cell antigen), and *ompB* (outer membrane protein B). Detection rates ranged from 86.7% to 100% in females but below 20% in both male groups for six of seven genes (Fig. 1c). Quantitative analysis confirmed significantly higher expression in females across all genes, with male levels at or near baseline regardless of mating status (Fig. 1d-j). To confirm this was not due to low bacterial load, we measured *gltA* RNA/DNA ratios in DNA-positive males and DNA-positive females. Because *gltA* is present as a single-copy gene in the *Rb* genome, the RNA/DNA ratio provides an estimate of transcriptional activity per bacterium. RNA/DNA ratios were significantly lower in males than females (Fig. S0), indicating reduced per-bacterium transcriptional activity rather than a low-load artifact. These results show that although *Rb* DNA is detectable in male *I. scapularis*, the bacterium is transcriptionally inactive. Since *Rb* relies exclusively on transovarial transmission, male ticks represent a reproductive dead end for this endosymbiont.

**Fig. 1.**
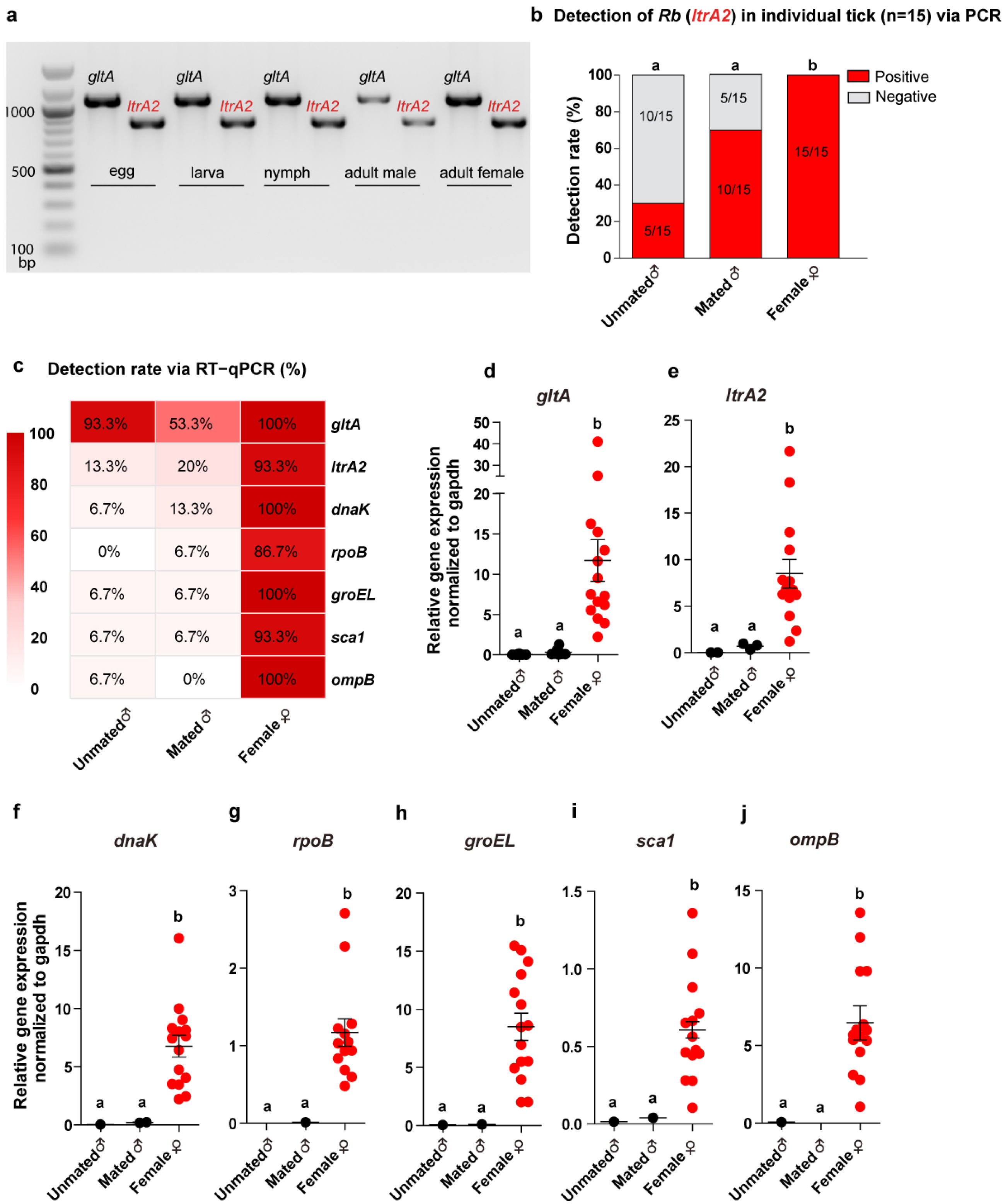
*Rb* is present across *I. scapularis* life stages but transcriptionally silent in male ticks. (**a**) PCR detection of *Rb* using *gltA* and *ltrA2* primers across tick developmental stages (egg, larva, nymph, adult male, and adult female). (**b**) Proportion of *Rb*-positive (*ltrA2*) individuals among unmated males, mated males, and females (n = 15 per group). Red, positive; gray, negative. Numbers indicate positive/total. Groups with different letters are significantly different (Fisher’s exact test with Bonferroni correction, p < 0.05) (**c**) Heatmap of RT-qPCR detection rates (%) for seven *Rb* genes across the three groups. Color scale: white (0%) to red (100%). (**d-j**) Relative expression of *gltA* (**d**), *ltrA2* (**e**), *dnaK* (**f**), *rpoB* (**g**), *groEL* (**h**), *sca1* (**i**), and *ompB* (**j**) normalized to tick gapdh. Each dot represents one individual tick (n = 15 per group). Bars indicate mean ± SEM. Groups with different letters are significantly different (Kruskal-Wallis test with Dunn’s post hoc correction, p < 0.05).

### *Rb* is exclusively localized within the reproductive tissue of unfed female *I. scapularis*

We next examined adult females, where *Rb* is transcriptionally active and responsible for transovarial transmission. Although *Rb* DNA was detected in larvae and nymphs (Fig. 1a), we focused on adult females given their central role in *Rb* maintenance and vertical transmission. The tissue distribution of *Rb* within female ticks is directly relevant to its biological function, yet remains debated, as some studies have suggested that *Rb* may also colonize salivary glands (18). To resolve this question, we examined dissected ovary, salivary gland, and midgut tissues from the same cohort of unfed female ticks using three complementary approaches: TEM, DNA FISH, and tissue-specific PCR.

Low-magnification TEM of ovary tissues revealed oocytes at multiple developmental stages (oo I-III), interstitial tissue cells (ITC), the oviduct (ov), and surrounding hemolymph cells (HMC) (Fig. 2a). At higher magnification, *Rickettsia*-like organisms were frequently observed and often densely distributed within stage I and stage II oocytes (oo I, oo II) as well as in interstitial cells adjacent to the oviduct (od) (Fig. 2b, red arrows). *Rb* was also observed in stage III oocytes (oo III), where bacteria were present in the cytoplasm near the nucleus (n) (Fig. 2c). The relatively high density of bacteria observed across multiple oocyte stages indicates that *Rb* is abundant in ovaries of unfed female ticks. Within interstitial cells, *Rb* appeared in autophagic vacuoles (AVd) as well (Fig. 2d), suggesting potential interactions between the endosymbiont and host vesicular compartments. In contrast, the pedicel region, containing epithelial pedicel cells (EPC), pigment granules (pg), mitochondria (mi), and nearby hemolymph cells (HMC), lacked any detectable *Rickettsia*-like structures (Fig. 2e), suggesting that *Rb* does not broadly colonize non-reproductive support cells and was not observed in adjacent hemolymph-associated regions. To further confirm TEM observations, DNA FISH using an ltrA2-specific probe was performed on tissues from the same tick cohort. In ovary sections, distinct *Rb*-specific signals (red) were detected within oocyte-containing regions (Fig. 2f). Higher-magnification views of two independent regions of interest confirmed discrete bacterial signals in both areas (Fig. 2f, panels 1-4), with no signal detected in salivary gland or midgut tissues (Fig. 2g-h). To further examine tissue distribution across individuals, PCR targeting *gltA* and *ltrA2* was performed on dissected tissues from 20 unfed female ticks. *Rb* DNA was detected in 100% (20/20) of ovary samples using gltA and 95% (19/20) using *ltrA2*, while all salivary gland and midgut samples were negative for both markers (0/20) (Fig. 2i).

**Fig. 2.**
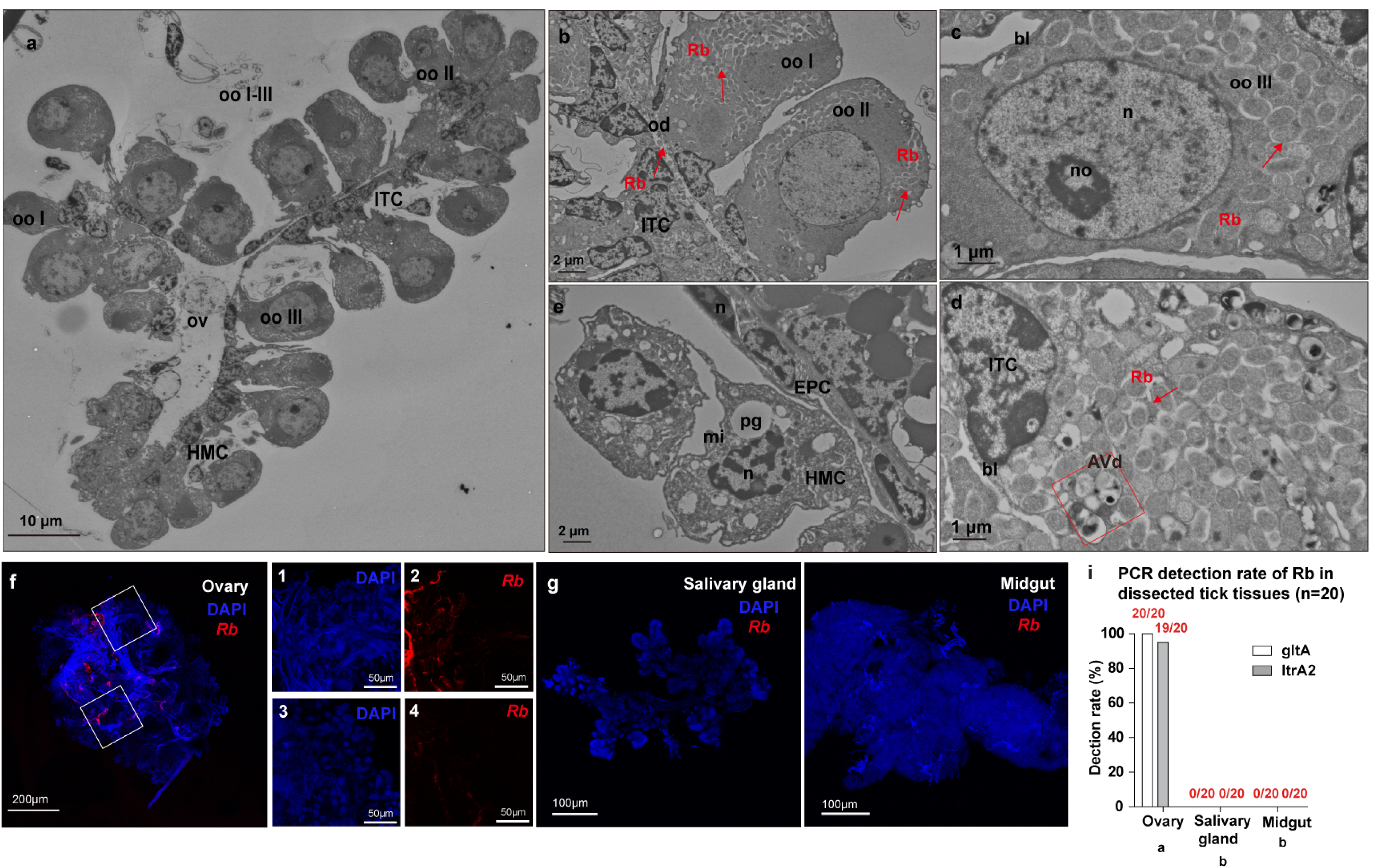
*Rb* is restricted to ovarian tissues in unfed female *I. scapularis*. (**a-e**) Transmission electron microscopy (TEM) of dissected ovary tissues from unfed females. (**a**) Low-magnification overview showing oocytes at multiple developmental stages (oo I-III), interstitial tissue cells (ITC), oviduct (ov), and hemolymph cells (HMC). (**b**) Higher magnification view showing *Rb* (red arrows) within stage I and II oocytes (oo I, oo II) and interstitial tissue cells (ITC) adjacent to the oviduct (od). (**c**) *Rb* within a stage III oocyte (oo III), distributed in the cytoplasm adjacent to the nucleus (n) and nucleolus (no). Basal lamina (bl). (**d**) *Rb* observed within interstitial tissue cells (ITC) adjacent to an autophagic vacuole (AVd). (**e**) Pedicel region containing epithelial pedicel cells (EPC), pigment granules (pg), mitochondria (mi), and hemolymph cells (HMC), with no detectable *Rb*. (**f**) DNA FISH using an *ltrA2*-specific probe (red) on ovary tissue sections; tick cell nuclei counterstained with DAPI (blue). Panels 1-4 show higher-magnification views of the two boxed regions. Panels 1 and 3 show DAPI signal; panels 2 and 4 show the corresponding *Rb* signal. (**g-h**) DNA FISH of salivary gland (**g**) and midgut (**h**) tissues showing no detectable *Rb* signal. (**i**) PCR detection rates using *gltA* and *ltrA2* primers in dissected tissues from unfed females (n = 20 per tissue). Tissues with different letters show significantly different detection rates (Fisher’s exact test, p < 0.001). Numbers indicate positive/total samples tested.

To further assess whether *Rb* colonizes non-reproductive tissues, TEM was performed on dissected salivary glands and midgut from the same unfed female cohort. Salivary gland examination included all three major acinar cell types (types I-III) (Supplementary Fig. S1a-d), and midgut analysis included epithelial cells (Supplementary Fig. S2a-b) along the entire length of the tissue. No Rickettsia-like organisms were detected in either tissue. Taken together, these results demonstrate that *Rb* in unfed female *I. scapularis* is restricted to ovarian tissues and absent from salivary glands, midgut, and hemolymph.

### Reduced density of *Rb* in ovaries of engorged female ticks

After blood feeding, female *I. scapularis* undergo a major physiological transition characterized by vitellogenesis and ovarian maturation (19). Because *Rb* was abundant in ovaries of unfed females (Fig. 2), we next examined whether symbiont abundance changes following engorgement. Semi-thin sections confirmed extensive ovarian development in engorged females, with oocytes spanning multiple developmental stages (oo I-V) and progressive yolk deposition (Fig. 3a). TEM revealed oocyte cytoplasmic expansion and yolk platelet (yp) development during vitellogenesis (Fig. 3b-f).

**Fig. 3.**
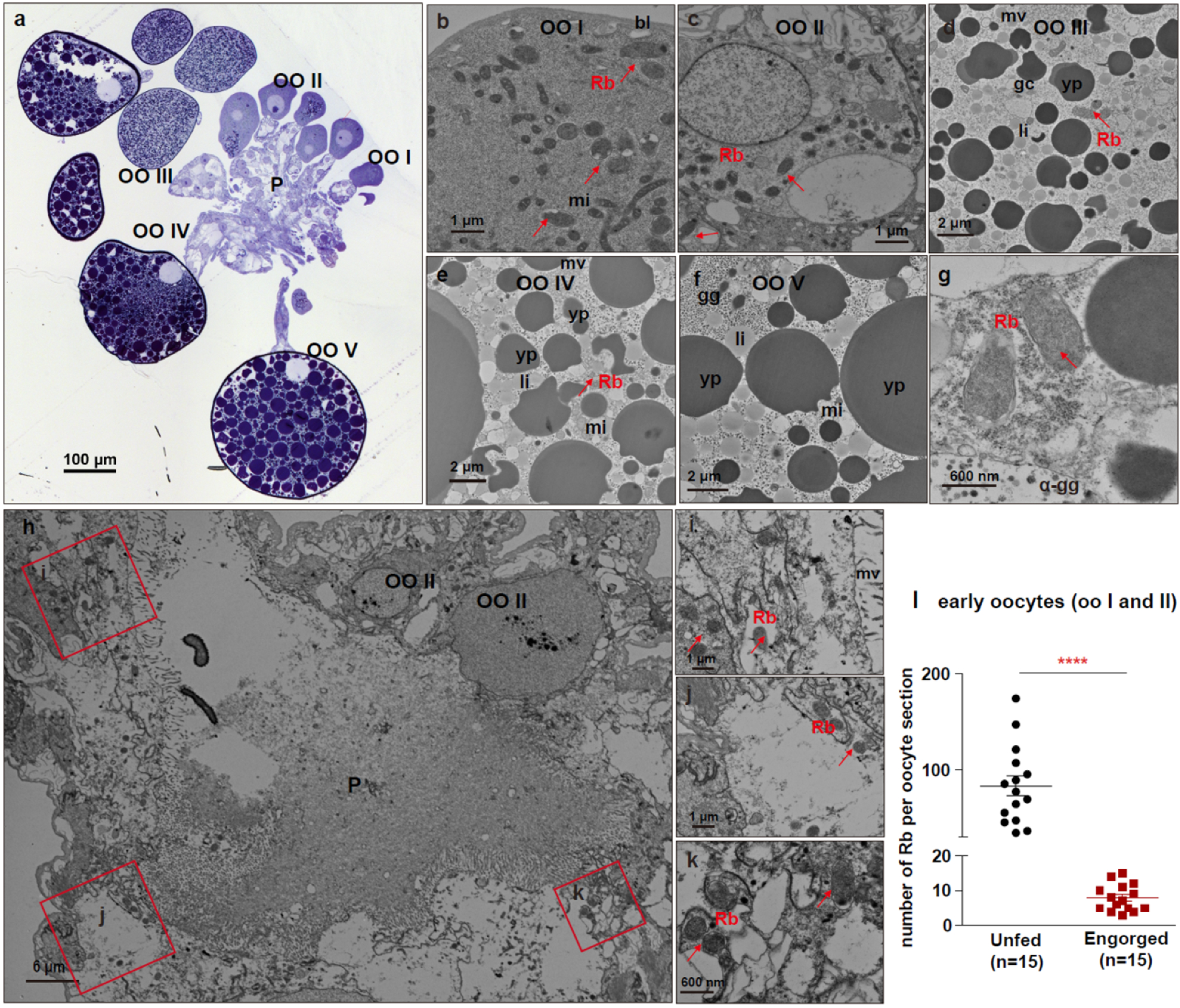
*Rb* abundance is markedly reduced in ovaries of engorged female *I. scapularis*. (**a**) Toluidine blue-stained semithin section of a resin-embedded ovary from a fully engorged female, showing oocytes at progressive developmental stages (oo I-V) and the pedicel region (P). (**b-f**) TEM images of oocytes at successive developmental stages from engorged females: oo I (**b**), oo II (**c**), oo III (**d**), oo IV (**e**), and oo V (**f**). Progressive maturation is accompanied by cytoplasmic enlargement and yolk platelet (yp) accumulation. Rare *Rb* are indicated by red arrows. (**g**) Higher-magnification TEM image showing representative *Rb* within late-stage oocyte cytoplasm. (**h**) Low-magnification TEM overview of the pedicel region (P) in engorged ovaries, showing extensive structural degradation. (**i-k**) Higher-magnification views of regions indicated in (**h**), showing rare *Rb* restricted to oocytes adjacent to the degraded pedicel. (**l**) Quantification of *Rb* cells per oocyte section in early-stage oocytes (oo I-II) from unfed and fully engorged females. Three ticks per group; five oo I-II sections per tick (n = 15 sections per group). Each dot represents one oocyte section. Bars indicate mean ± SEM. Statistical comparison performed on tick-level means (n = 3 per group) using a two-tailed Mann-Whitney U test (**** *P* < 0.0001). Abbreviations: oo, oocyte; P, pedicel; yp, yolk platelet; mi, mitochondria; mv, microvilli; bl, basal lamina; gc, Golgi complex; li, lipid droplet, gg, glycogen granules. α-gg, aggregated glycogen granules.

In contrast to the abundant *Rb* observed in ovaries of unfed females (Fig. 2), only rare *Rb* were detected in ovaries from engorged females. Occasional bacteria were observed within early oocytes (oo I-II) and in some vitellogenic oocytes (oo III-IV), but the density of *Rb* within developing oocytes appeared reduced relative to unfed females (Fig. 3b-e, red arrows). In late-stage oocytes containing large yolk platelets (oo V), *Rb* was rarely detected (Fig. 3f). Representative examples required systematic examination of multiple TEM sections (Fig. 3g). TEM examination also revealed substantial structural remodeling of the pedicel region in engorged ovaries. The pedicel region displayed extensive structural degradation and lacked detectable *Rb*; the few bacteria observed were restricted to adjacent oocytes (Fig. 3h-k).

To rule out dilution effects from oocyte enlargement during vitellogenesis, we performed TEM-based enumeration of *Rb* in size-matched early-stage oocytes (oo I-II) from three unfed and three fully engorged females (five sections per tick; n = 15 sections per group). Quantitative analysis confirmed that *Rb* density was significantly reduced in early oocytes of engorged females compared to unfed samples (Fig. 3l).

This comparison controlled for oocyte size and developmental stage. We note that these section-level counts reflect local *Rb* density within early oocytes and cannot be converted into the total number of *Rb* per oocyte or per egg; absolute loads at the whole-egg stage have been reported previously and are considered in the Discussion. Because salivary glands undergo programmed degeneration and the midgut becomes filled with blood meal material following engorgement (Supplementary Figs. S1e-f, S2c-e), isolating intact tissues for PCR or DNA FISH was not feasible. These results show that blood feeding and vitellogenesis are accompanied by marked structural remodeling of the ovary and a significant reduction in *Rb* density within developing oocytes.

### Characterization of the vitellogenesis gene repertoire identifies candidate regulators

We next examined the vitellogenesis pathway in *I. scapularis* to investigate its potential role in *Rb* regulation (Fig. 4a). Since *Rb* localizes within developing oocytes and becomes depleted following blood feeding (Figs. 2-3), we hypothesized that vitellogenesis, which is strongly induced by blood feeding, may regulate *Rb* dynamics during oocyte maturation. Genomic characterization identified 20 putative Vgs (vitellogenin synthesis) genes and 15 Vgr (vitellogenin receptor) genes in *I. scapularis*. Phylogenetic analysis revealed distinct evolutionary patterns between the two gene families. Vgs paralogs formed multiple well-supported sister clades and remained comparatively conserved, whereas Vgr paralogs exhibited substantially greater sequence divergence (Fig. 4b). Cross-species comparisons confirmed that Vgr diversification is a conserved feature of ticks, while Vgs copy numbers remain more constrained (Supplementary Figs. S3-S4; Table 1). Comparison with *D. melanogaster* revealed a pronounced tick-specific Vgr expansion relative to insects, further supported by broader analyses across arthropods, insects, and nematodes (Supplementary Figs. S5-S7; Table 1).

**Fig. 4.**
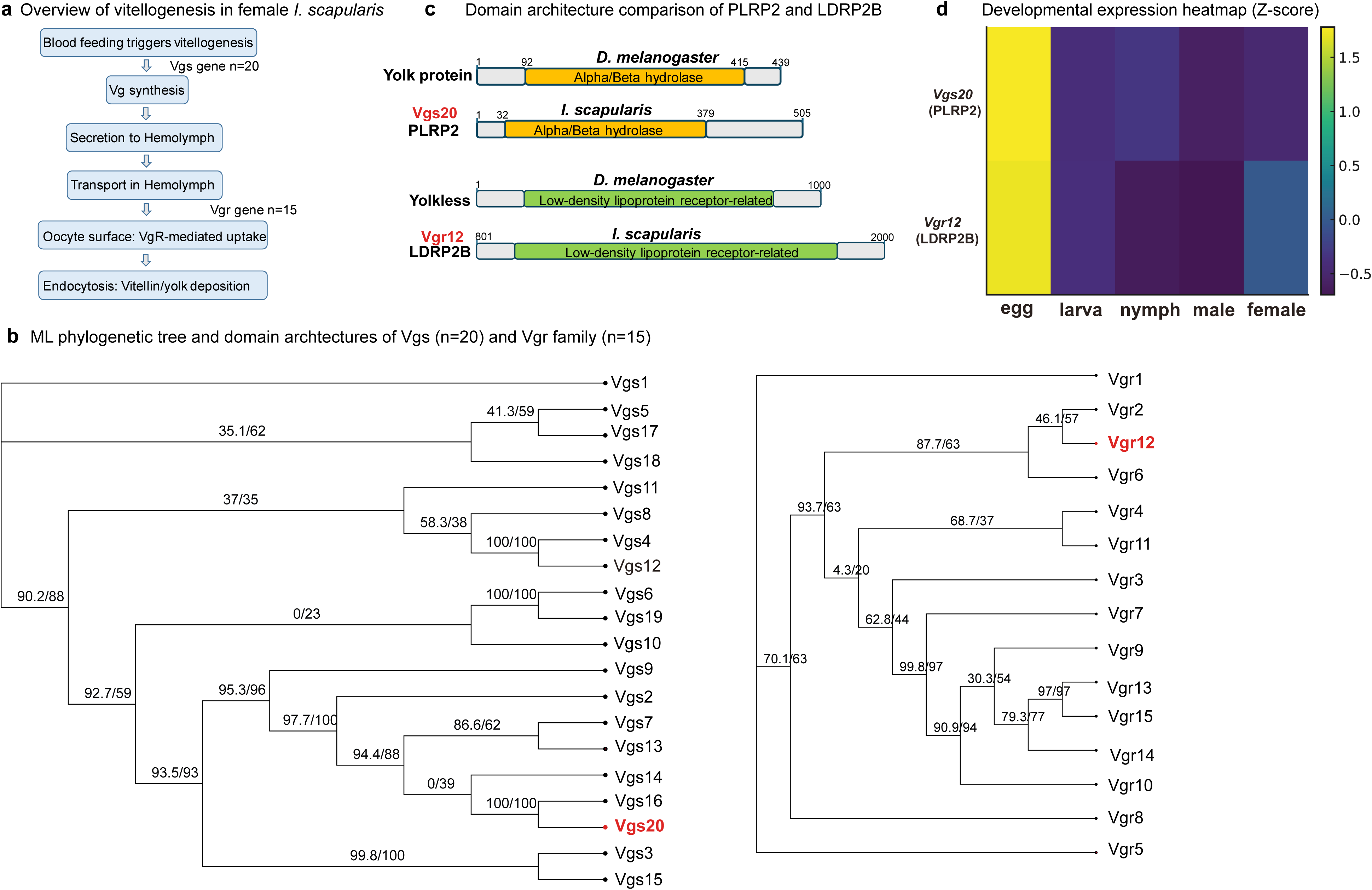
Genomic characterization of the vitellogenesis genes in *I. scapularis*. (**a**) Schematic of the vitellogenesis pathway in female ticks. Blood feeding triggers vitellogenin synthesis in peripheral tissues (Vgs, n = 20), secretion into the hemolymph and receptor-mediated uptake into developing oocytes via vitellogenin receptors (Vgr, n = 15). (**b**) Maximum likelihood (ML) phylogenetic trees of the Vgs family (left, n = 20) and Vgr family (right, n = 15). Branch support values are shown as SH-aLRT/UFBoot. Vgs proteins form tightly clustered clades indicating strong conservation; Vgr proteins show substantial phylogenetic divergence. Candidate genes Vgs20 and Vgr12 highlighted in red. (**c**) Domain architectures of Vgs20 (PLRP2) and Vgr12 (LDRP2B) compared with *D. melanogaster* homologs (yolk protein and yolkless receptor). Domain annotations based on InterPro/Pfam searches. Protein lengths (aa) indicated at right. (**d**) Developmental expression heatmap of Vgs20 and Vgr12 across five life stages (egg, larva, nymph, adult male, adult female), Relative expression was quantified by RT-qPCR, normalized to the tick housekeeping gene *gapdh*, and displayed as Z-scores. Dashed line separates the two genes.

**Table 1.**
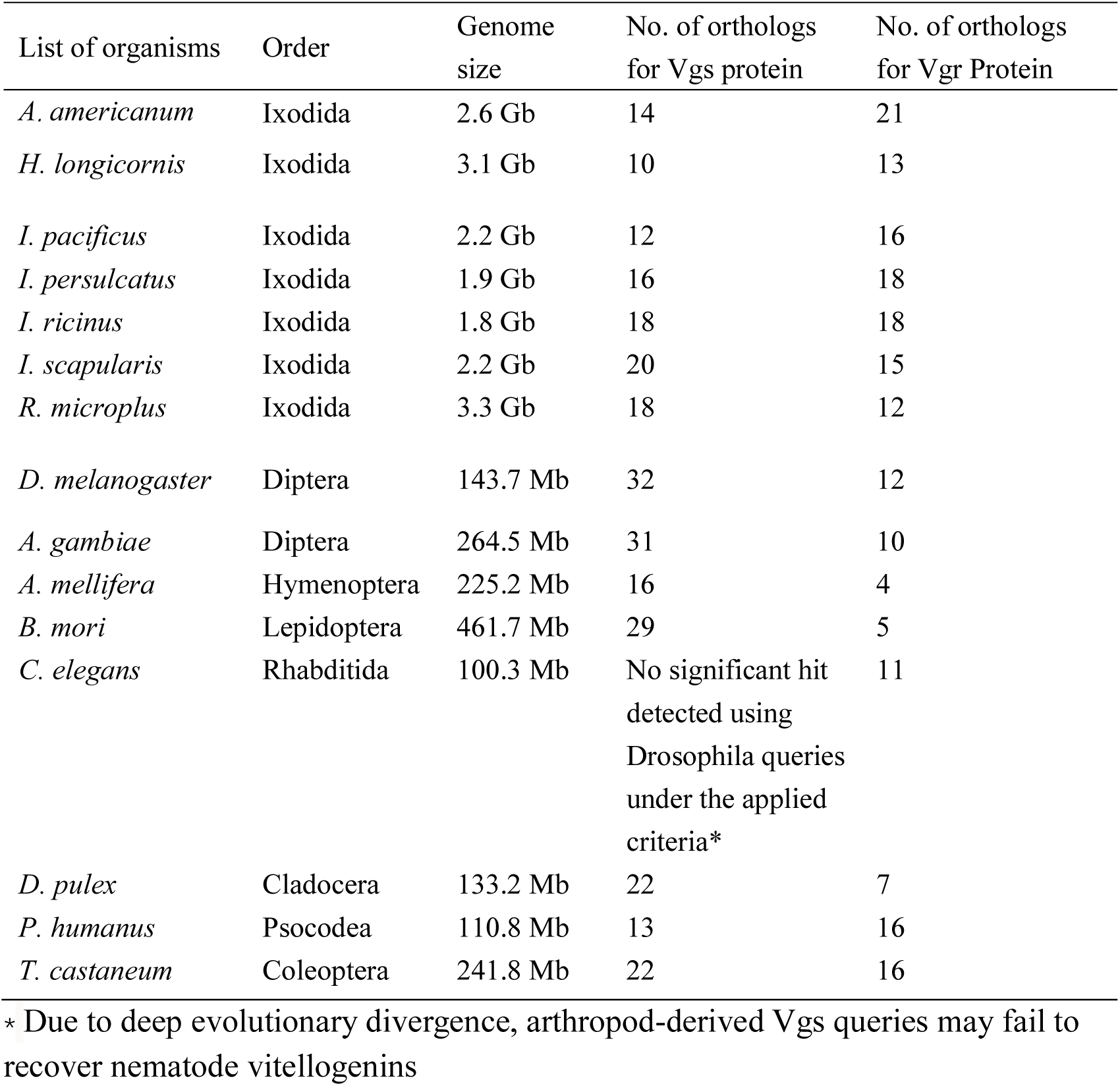
Comparative analysis of vitellogenin synthesis (Vgs) and vitellogenin receptor (Vgr) gene repertoires across arthropods and nematodes. The number of predicted orthologs for Vgs and Vgr proteins was identified from publicly available genome assemblies of representative species spanning Ixodida (ticks), Diptera, Hymenoptera, Lepidoptera, Coleoptera, Cladocera, Psocodea, and Rhabditida. Gene copy numbers identified from publicly available genome assemblies using BLAST searches with *I. scapularis* and *D. melanogaster* sequences as queries. Genome sizes were obtained from reference genome annotations. “No ortholog found” indicates that no significant homolog of Vg proteins was detected in the corresponding genome under the applied search criteria. This table highlights lineage-specific expansion or contraction of Vgs and Vgr gene families, particularly in tick species, suggesting adaptive significance in blood-feeding ecology.

Functional annotation indicated that most Vgs proteins are predicted to be extracellular, consistent with hemolymph secretion, whereas Vgr proteins displayed more subcellular localizations including the plasma membrane and endocytic compartments (Tables 2-3). Protein domain analysis further highlighted these differences between the two families. Vgs proteins predominantly contained conserved pancreatic lipase-like domains, whereas Vgr proteins displayed more diverse domain architectures, including low-density lipoprotein receptor (LDLR)-related, transmembrane, and endocytic domains, consistent with functional specialization (Fig. S9). Gene structure and conserved motif analyses further supported the overall distinction between the two families (Figs. S8-S10).

**Table 2.**
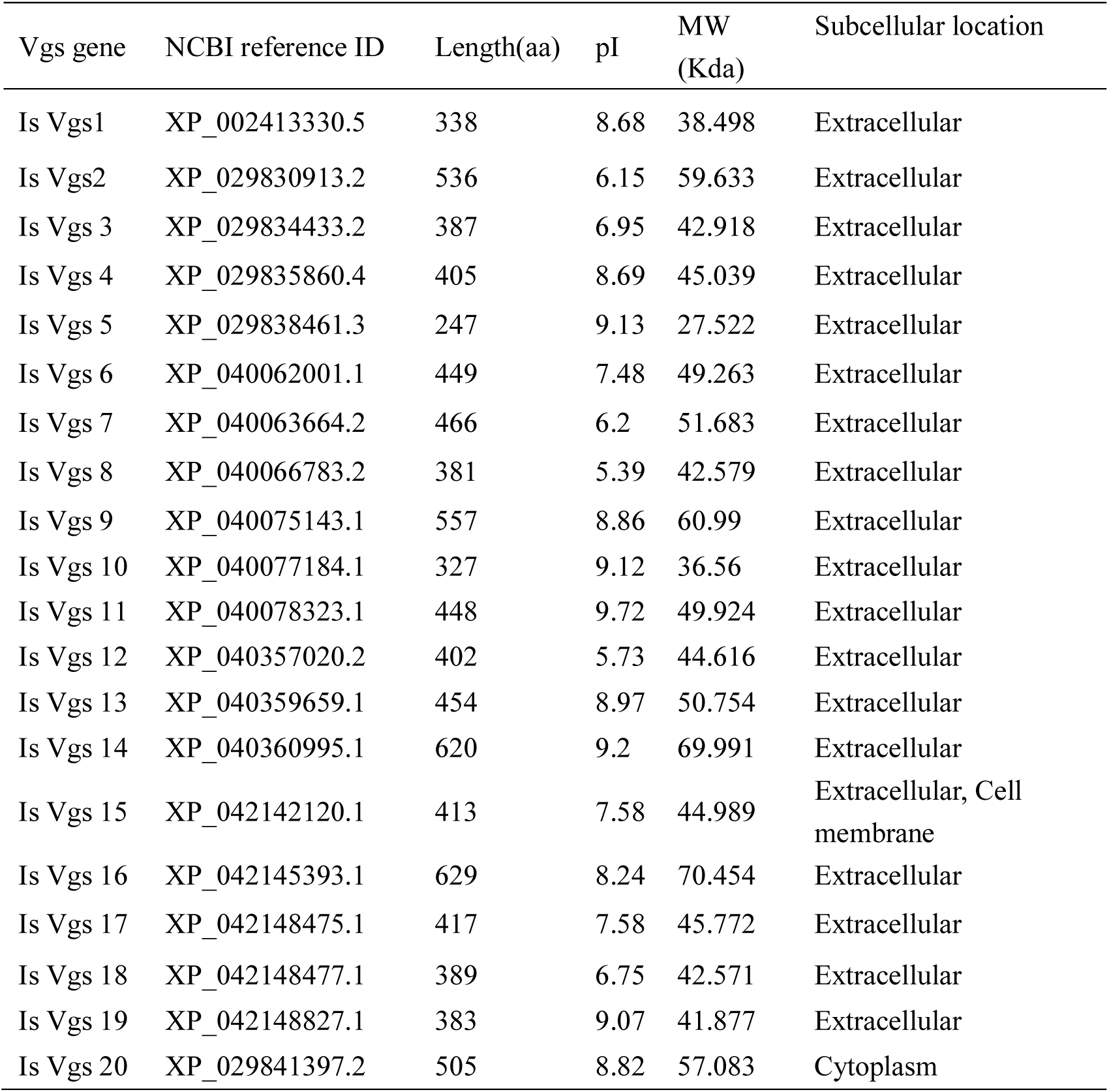
Predicted characteristics of putative vitellogenin synthesis (Vgs) proteins in *I. scapularis*. A total of 20 putative Vgs proteins were identified from the *I. scapularis* genome based on sequence homology. Protein length (amino acids), theoretical isoelectric point (pI), and molecular weight (MW) were predicted using standard bioinformatic tools. Subcellular localization predicted using DeepLoc 2.1 algorithm. Most Vgs proteins are predicted to be extracellular, consistent with their role as yolk protein precursors, while a small subset shows alternative localization patterns, suggesting potential functional diversification. NCBI accession numbers provided for sequence retrieval.

**Table 3.** Predicted characteristics of putative vitellogenin receptor (Vgr) proteins in *I. scapularis*. A total of 15 putative Vgr proteins were identified from the *I. scapularis* genome. Protein length, theoretical isoelectric point (pI), and molecular weight (MW) were predicted. Subcellular localization analysis indicates that most Vgr proteins are associated with the cell membrane, consistent with their function as receptors mediating vitellogenin uptake. Several Vgr proteins are additionally predicted to localize to lysosomal/vacuolar compartments or extracellular regions, suggesting possible variation in receptor trafficking or function. Size variation is pronounced (438-6336 aa), indicating potential functional specialization within the gene family. Vgr12 represents one of the largest family members with extensive LDLR-related domains.

| Vgr genes | NCBI reference ID | Length(aa) | pI | MW (Kda) | Subcellular location |
| --- | --- | --- | --- | --- | --- |
| Is Vgr 1 | XP_029827184.2 | 2129 | 5.59 | 231.658 | Cell membrane |
| Is Vgr 2 | XP_029830175.1 | 1550 | 6.02 | 170.917 | Cell membrane,<br>Lysosome/Vacuole |
| Is Vgr 3 | XP_029830196.2 | 1805 | 5.05 | 200.468 | Cell membrane,<br>Lysosome/Vacuole |
| Is Vgr 4 | XP_029831536.4 | 1857 | 5.38 | 206.425 | Cell membrane |
| Is Vgr 5 | XP_029834260.2 | 1124 | 6.48 | 124.756 | Cell membrane,<br>Lysosome/Vacuole |
| Is Vgr 6 | XP_029839443.3 | 882 | 4.84 | 98.075 | Cell membrane,<br>Lysosome/Vacuole |
| Is Vgr 7 | XP_029843234.2 | 1266 | 5.51 | 136.148 | Extracellular |
| Is Vgr 8 | XP_029850375.2 | 4668 | 5.14 | 515.924 | Cell membrane |
| Is Vgr 9 | XP_029851566.2 | 1211 | 5.17 | 133.857 | Extracellular |
| Is Vgr 10 | XP_040073621.1 | 927 | 6.98 | 101.078 | Extracellular |
| Is Vgr 11 | XP_040078195.1 | 4410 | 5.32 | 483.263 | Extracellular |
| Is Vgr 12 | XP_042143231.1 | 4583 | 5.21 | 512.859 | Cell membrane |
| Is Vgr 13 | XP_042144874.1 | 2945 | 5.07 | 318.518 | Extracellular |
| Is Vgr 14 | XP_042145593.1 | 438 | 4.11 | 48.822 | Cytoplasm,<br>Extracellular |
| Is Vgr 15 | XP_042146222.1 | 6336 | 5.51 | 686.715 | Cell membrane |

We prioritized candidates using phylogenetic position (divergent placement relative to the majority of family members), domain architecture, predicted subcellular localization, and developmental expression profiles. Vgs20 (PLRP2) and Vgr12 (LDRP2B) emerged as top candidates. Vgs20 contains a pancreatic lipase-like domain similar to *D. melanogaster* yolk protein but is distinguished by an atypical predicted cytoplasmic localization relative to other Vgs members, whereas Vgr12 harbors extensive LDLR-related domains and transmembrane regions comparable to *D. melanogaster* yolkless receptor and consistent with receptor-mediated endocytosis (Fig. 4c). Both genes show peak expression at the egg stage and reduced expression during larval and nymph stages; Vgr12 displays a marked female-specific re-induction in adults, consistent with vitellogenesis activation during oocyte maturation (Fig. 4d). Together, these observations identified Vgs20 and Vgr12 as candidates for subsequent functional analysis.

To further evaluate the evolutionary relationship between the Vgs and Vgr gene families, we compared sequence conservation using pairwise amino acid identity (Fig. 5). Pairwise identity was higher among Vgs paralogs than among Vgr paralogs (median identity, 0.21 vs. 0.06; Fig. 5a). Mean sequence identity was also significantly higher in the Vgs family than in the Vgr family (0.24 ± 0.05 vs. 0.05 ± 0.03; Mann-Whitney U test, P = 6.2 × 10⁻¹⁰; Fig. 5b). Together, these results show that the Vgs family is more conserved, whereas the Vgr family has undergone greater sequence diversification.

**Fig. 5.**
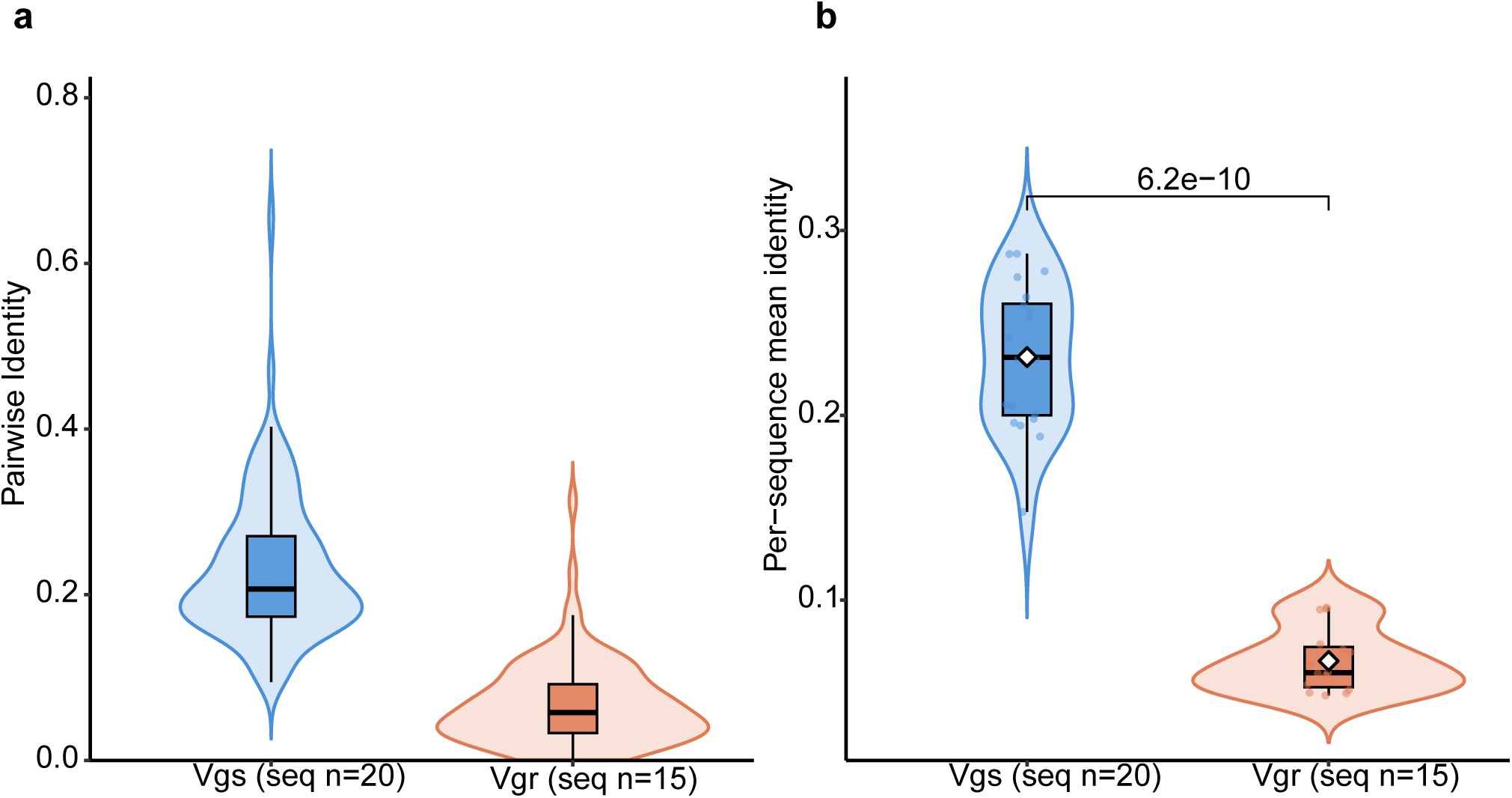
Quantitative comparison of sequence conservation between the Vgs and Vgr gene families. (**a**) Distribution of pairwise amino acid sequence identity among all within-family comparisons for Vgs (20 sequences; 190 pairwise comparisons) and Vgr (15 sequences; 105 pairwise comparisons). Pairwise identity was calculated as the proportion of identical amino acid residues over aligned positions, excluding gap-gap positions. (**b**) Mean sequence identity for each Vgs and Vgr protein, calculated by averaging its pairwise identity values against all other members of the same gene family. Horizontal lines indicate the median. Statistical significance was determined using a two-sided Mann-Whitney U test.

### *In vitro* and *in vivo* validation identifies Vgs20 and Vgr12 as differential regulators of *Rb* load and transcriptional state

We functionally validated the roles of Vgs20 (PLRP2) and Vgr12 (LDRP2B) in regulating *Rb* using complementary in vitro and in vivo systems. In ISE6 tick cells, *Rb* infection significantly suppressed expression of both *Vgs20* and *Vgr12* (Fig. 6a). RNAi-mediated silencing achieved partial knockdown of both targets (Fig. 6b) and significantly reduced *Rb* genomic copies (Fig. 6c). We extended these findings to unfed adult female ticks, prior to blood feeding and active vitellogenesis. *Rb*-infected ticks showed significantly lower expression of both *Vgs20* and *Vgr12* compared to *Rb*-free ticks (Fig. 6d), confirming that *Rb* suppresses this pathway before reproductive maturation begins. RNAi silencing was more efficient *in vivo* (Fig. 6e). However, the impact on bacterial load diverged: *Vgr12* silencing significantly reduced *Rb* genomic copies, while *Vgs20* silencing produced no significant change (Fig. 6f). This suggests that *Vgr12* directly supports bacterial maintenance, while *Vgs20* influences *Rb* through a load-independent mechanism.

**Fig. 6.**
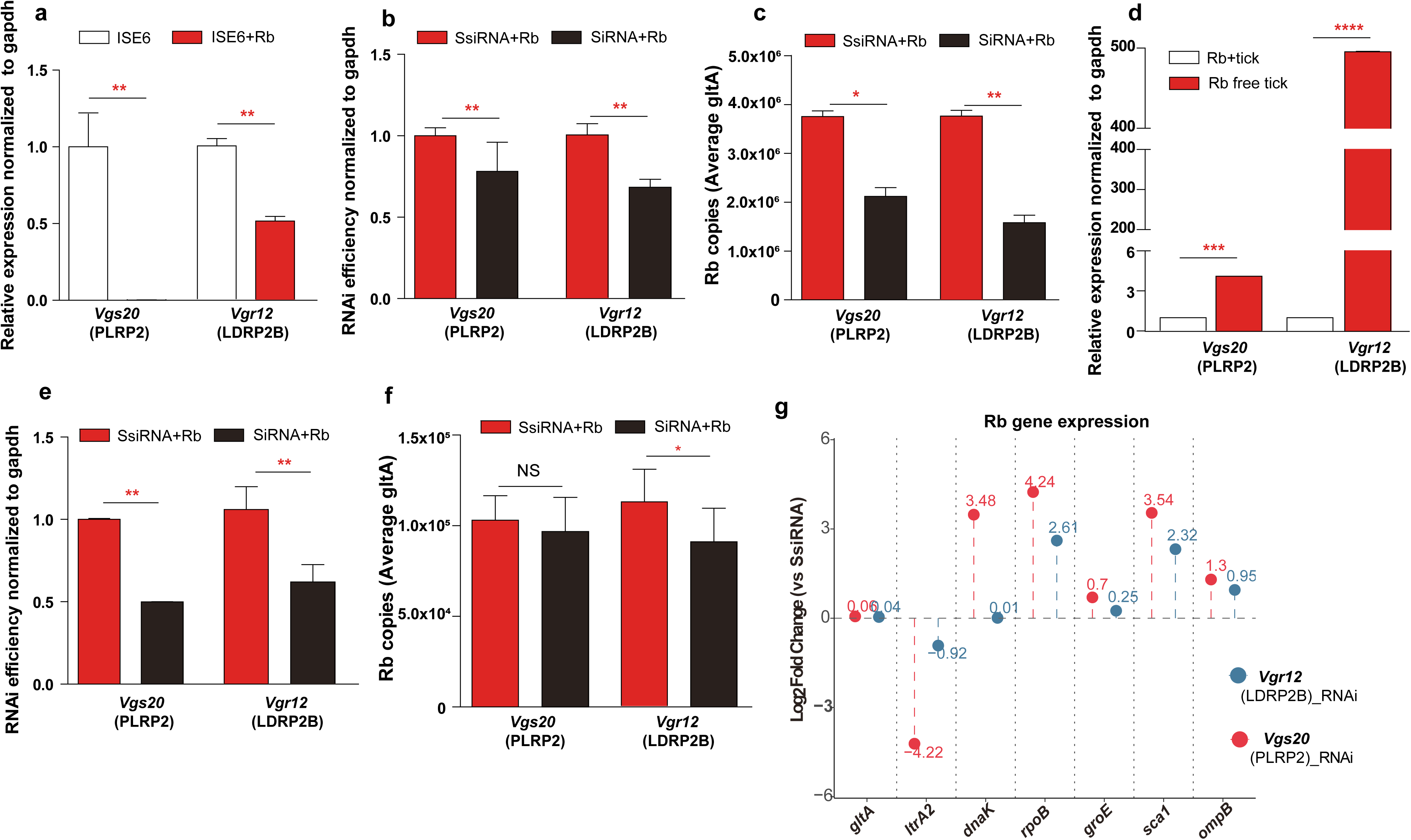
Vgs20 and Vgr12 differentially regulate *Rb* load and transcriptional state. (**a**) Relative expression of Vgs20 and Vgr12 in ISE6 tick cells infected with *Rb* compared to uninfected controls, normalized to *gapdh*. (**b**) RNAi silencing efficiency of Vgs20 and Vgr12 in ISE6 cells, normalized to *gapdh*. (**c**) *Rb* genomic copy number following RNAi silencing in ISE6 cells, quantified by qPCR targeting *gltA*. (**d**) Relative expression of *Vgs20* and *Vgr12* in *Rb*-infected versus *Rb*-free unfed adult female ticks, normalized to *gapdh*. (**e**) RNAi silencing efficiency of *Vgs20* and *Vgr12* in unfed female ticks, normalized to *gapdh*. (**f**) *Rb* genomic copy number in unfed ticks following RNAi treatment, quantified by qPCR targeting *gltA*. *Vgr12* silencing significantly reduced *Rb* load; *Vgs20* silencing did not produce a significant change at this stage. (**g**) Transcriptional response of *Rb* to Vgs20 and Vgr12 silencing in unfed ticks. Log2 fold change of seven *Rb* genes relative to SsiRNA controls is shown for Vgr12 RNAi (blue) and Vgs20 RNAi (red). Values above or below each point indicate Log2 fold change. For all panels, data represent mean ± SEM. Statistical significance was determined using a two-tailed Student’s t-test. NS, not significant; *P < 0.05; **P < 0.01; ***P < 0.001; ****P < 0.0001.

Both silencing treatments markedly altered *Rb* transcription regardless of bacterial load. Expression of seven *Rb* genes representing distinct functional categories was quantified as Log2 fold change relative to SsiRNA controls (Fig. 6g). The housekeeping gene *gltA* showed no significant change in either group (Log_2_ FC ≈ 0), consistent with stable basal metabolic activity per bacterium. In contrast, the *Rb*-specific plasmid marker *ltrA2* was strongly downregulated in both groups (Log_2_ FC < −4). Stress response, transcription, and surface presentation genes, *dnaK, rpoB, groEL, sca1*, and *ompB*, were significantly upregulated in both RNAi groups, with several exceeding Log₂ FC > 3 (Fig. 6g). Importantly, even when bacterial load remained unchanged (*Vgs20* group), the symbiont population shifted toward a stress-responsive transcriptional state. These results show that the host vitellogenesis pathway maintains *Rb* transcriptional quiescence, and that disruption of this regulation activates symbiont stress responses regardless of bacterial load.

## DISCUSSION

This study defines a working model for how *Rb* is regulated within the reproductive biology of its obligate tick host, *I. scapularis* (Fig. 7). Obligate, vertically transmitted endosymbionts must maintain a tight balance with host reproductive physiology: insufficient activity compromises transmission, whereas uncontrolled proliferation risks host fitness costs and symbiont clearance (20–21). Our findings reveal that this balance is enforced through three coordinated layers: (1) sex-dependent transcriptional silencing that restricts active *Rb* to females, (2) strict ovary-specific localization, and (3) a vitellogenesis-associated regulatory axis that maintains *Rb* in a quiescent state during oocyte maturation. We also demonstrate bidirectional regulation: disruption of vitellogenesis alters *Rb* dynamics, whereas depletion of *Rb* correspondingly affects Vgs/Vgr expression (Fig. 7). These findings show that tick vitellogenesis functions as a key regulatory interface governing endosymbiont persistence during transovarial transmission, beyond its role in nutrient transport.

**Fig. 7.**
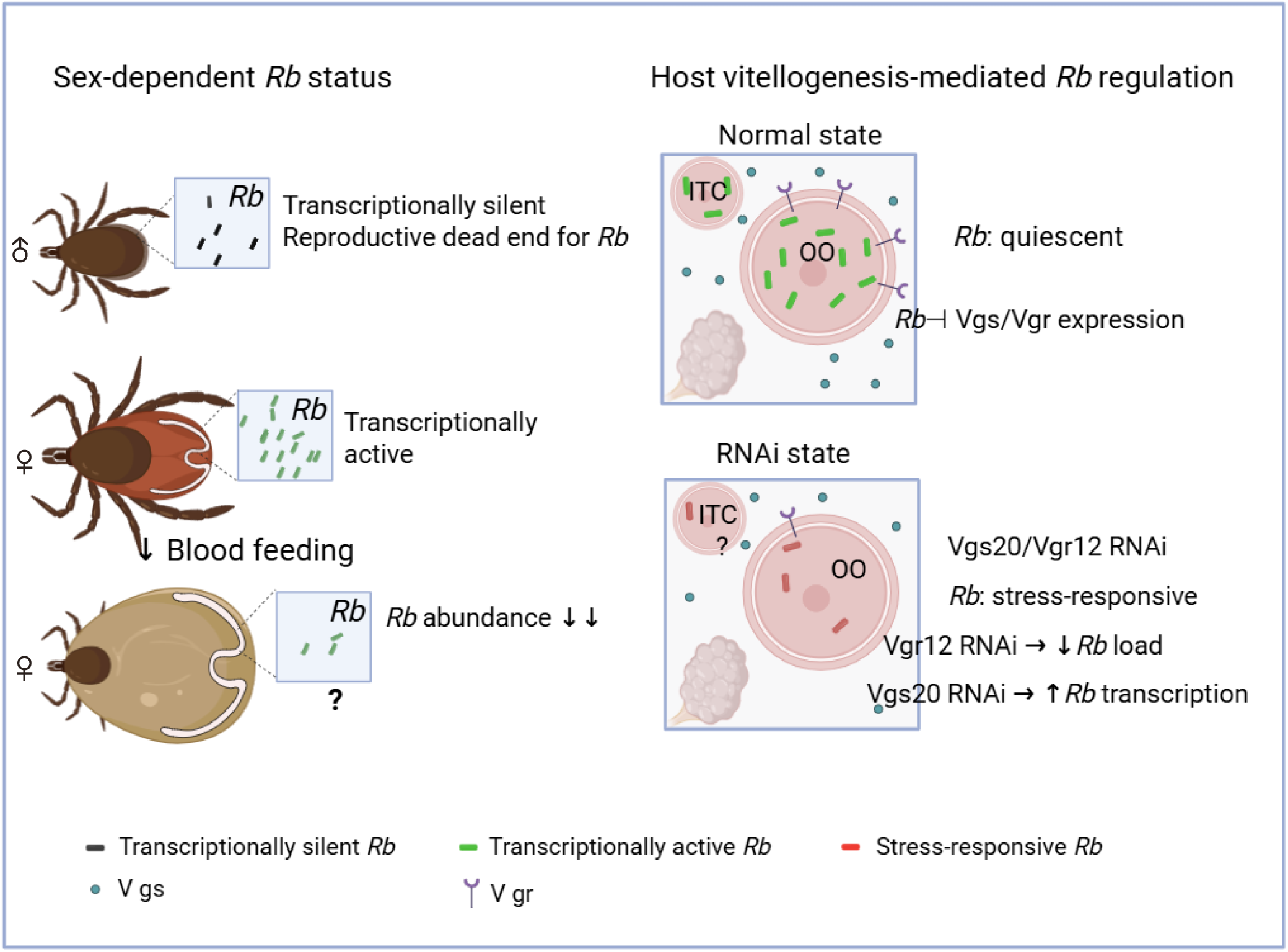
Proposed model for vitellogenesis-mediated regulation of *Rb* in *I. scapularis*. Left panel: sex-dependent *Rb* transcriptional status across tick physiological states. In males, *Rb* DNA is detectable but transcriptionally silent, representing a reproductive dead end for this obligate transovarially transmitted endosymbiont. In unfed females, *Rb* is transcriptionally active and restricted to ovarian tissues. Following blood feeding, *Rb* abundance is markedly reduced in engorged females; mechanism of depletion remains to be determined (?). Right panel: vitellogenesis-mediated regulation of *Rb* in unfed female ovarian tissue. Under normal conditions, *Rb* actively suppresses host Vgs20 and Vgr12 expression (Rb⊣Vgs/Vgr), while residual Vgs/Vgr activity maintains *Rb* transcriptionally quiescent. RNAi-mediated silencing of Vgs20 and Vgr12 disrupts this equilibrium: Vgr12 knockdown reduces *Rb* genomic load, whereas Vgs20 knockdown drives a shift toward a stress-responsive transcriptional state without immediately affecting bacterial numbers. ITC, interstitial tissue cell; OO, oocyte. Black bacteria, transcriptionally silent *Rb*; green bacteria, transcriptionally active quiescent *Rb*; red bacteria, stress-responsive *Rb* (RNAi-induced). Blue circles, Vg (vitellogenin); purple Y-shapes, Vgr (vitellogenin receptor).

*Rb* is transcriptionally quiescent in male *I. scapularis*. Although *Rb* DNA is detectable in males, expression of multiple *Rb* genes remains at or near baseline, and the reduced RNA/DNA ratio cannot be explained by a simple low-load artifact, instead supporting transcriptional repression on a per-bacterium basis (Fig. 1). While *Rb* DNA has been reported in males in previous field studies (10, 22), our data indicate that this does not represent a metabolically active symbiont population. This quiescence is consistent with the reproductive dead-end status of male ticks, which cannot contribute to transovarial transmission; maintaining metabolic activity in a non-transmitting host would provide limited fitness benefit at potential cost to the host (23).

Whether this silencing reflects active host regulation or passive metabolic downregulation remains unknown. The absence of vitellogenesis and the oocyte-associated niche in adult males suggests that host physiological environment, rather than immune suppression, is the primary determinant of symbiont transcriptional state. This contrasts with pathogenic spotted fever group rickettsiae, which have been detected in male ticks and can, in some cases, be sexually transmitted to females (24–26), highlighting a fundamental difference in transmission strategy between obligate endosymbionts and horizontally transmissible pathogens. How *Rb* behaves prior to adult sexual differentiation remains an open question. Although *Rb* DNA is detectable across immature stages (Fig. 1a), its transcriptional activity and tissue localization during larva and nymph development remain entirely uncharacterized. It will also be important to determine whether *Rb* abundance or transcription differs between immature ticks destined to develop into adult males or females. Whether *Rb* has additional stage-specific functions beyond reproduction, analogous to those described for endosymbionts such as *Wolbachia* in dipteran hosts (27–29), including potential nutritional roles suggested by its genome, remains to be determined.

The consistent restriction of *Rb* to ovarian tissues across multiple physiological states defines a second layer of host-symbiont specialization. *Rb* localizes to developing oocytes and interstitial ovarian cells, with no evidence of colonization in salivary glands or midgut in either unfed or engorged females. This consistent tissue tropism supports the view that *Rb* occupies a dedicated reproductive niche rather than a tissue-generalist distribution. In contrast to tick-borne pathogens, which colonize multiple tissues, including salivary glands, to enable horizontal transmission (30–31), *Rb* is confined to the ovary and reliant on transovarial transmission. The ovarian restriction parallels the germline tropism of *Wolbachia* in dipteran hosts (32, 2), suggesting convergent evolution toward reproductive tissue specialization among vertically transmitted arthropod endosymbionts.

*Rb* was frequently observed within autophagic compartments in ovarian interstitial cells (Fig. 2d), and our previous study in unfed female ticks demonstrated that *Rb* was associated with both autophagosomes and autolysosomes in ovarian tissues, and that autophagic compartments contained not only *Rb* but also cytoplasmic organelles and Golgi apparatus, a pattern consistent with non-selective autophagy rather than targeted xenophagy (14). These observations indicate that host autophagic activity constitutively engages *Rb* as part of a broader, non-selective intracellular remodeling process, contributing to baseline symbiont population control within ovarian tissues rather than targeted clearance. How this autophagic activity relates to the more dramatic remodeling triggered by blood feeding is a key question. Following engorgement, *Rb* density declined markedly in developing oocytes (Fig. 3), coinciding with the onset of receptor-mediated vitellogenin endocytosis (33–34) and large-scale intracellular remodeling during vitellogenesis. The temporal overlap between this surge in endocytic activity and *Rb* depletion suggests that blood-feeding-induced intracellular remodeling may contribute to *Rb* clearance beyond autophagic activity. Rather than requiring a dedicated symbiont-clearance mechanism, this reduction may emerge as a byproduct of the host’s broader reproductive physiology, showing a passive but consequential outcome of oocyte maturation, consistent with physiology-dependent modulation of endosymbiont density documented in other insect-symbiont systems (4). The decline of *Rb* in early oocytes reflects not just a consequence of oocyte expansion, but a co-adapted regulatory process that balances symbiont persistence with host reproductive demands (4). Our TEM counts quantify local *Rb* density within size-matched early oocytes and cannot be converted into total *Rb* per oocyte or per egg. Total symbiont abundance across oogenesis therefore remains unresolved: a net reduction during vitellogenesis, an approximately constant load with a transient decline in density, and an early reduction followed by recovery during late oogenesis are all compatible with our data. Whole eggs of *I. scapularis* have been reported to contain ∼5-10 × 10³ *Rb* (43), within the range expected for early oocytes, raising the possibility that symbiont numbers are re-established as oocytes mature. The persistence of rare *Rb* following engorgement is consistent with a transient controlled transmission bottleneck (35), suggesting that vitellogenesis may limit symbiont abundance while preserving enough bacteria for transovarial transmission.

The lineage-specific expansion of the Vgr gene family represents a significant evolutionary dimension of this study. Unlike insects, in which Vgr orthologs occupy single-copy positions within conserved pan-arthropod clades, *I. scapularis* encodes 15 distinct Vgr paralogs with substantially divergent domain architectures, while the Vgs family, despite comprising 20 members, displays only three principal domain configurations (Supplementary Figs. S4, S8-S10). This pattern, a conserved ligand family paired with a diversified receptor family, is consistent with the well-established evolutionary tendency for receptor proteins to undergo more extensive duplication and functional divergence than their ligands (36–37). We propose that blood-feeding-driven vitellogenin uptake created the selective pressure for Vgr expansion, and that this diversification reflects a conserved feature of the tick lineage rather than a species-specific anomaly (Supplementary Figs. S3-S4; Table 1). The expanded Vgr family suggests that receptor diversification has supported functions beyond vitellogenin uptake. Our data suggest that regulation of the endosymbiont *Rb* may be one such function. Whether this regulatory role contributed to the retention of specific Vgr paralogs, or evolved after receptor diversification for other biological functions, cannot be distinguished from the current data.

Consistent with theoretical predictions that receptor gene duplication facilitates neofunctionalization (36–37), our functional data suggest that specific Vgr paralogs have been co-opted for endosymbiont regulation. Vgs20 and Vgr12 emerged as top candidates based on atypical domain features and tightly coordinated temporal expression during oocyte maturation (Fig. 4 and 5). *Rb* actively suppresses both genes *in vitro* and *in vivo* (Fig. 6a, d), and RNAi-mediated knockdown triggers significant transcriptional reactivation of the surviving symbiont population (Fig. 6g). That *Rb* depletion restores Vgs/Vgr expression indicates that the symbiont actively modulates the host pathway governing its own transcriptional state. This suggests that the vitellogenesis machinery has, over evolutionary time, become a co-evolutionary relationship between host reproductive physiology and endosymbiont regulation (4).

A key functional outcome of this regulatory relationship is the decoupling of bacterial load from transcriptional state. Vgr12 silencing significantly reduced *Rb* genomic copies, whereas Vgs20 knockdown did not alter bacterial load but drove an equivalent transcriptional shift: *ltrA2* was strongly downregulated, while stress-response genes (*dnaK, groEL, rpoB*) and surface antigen genes (*sca1, ompB*) were significantly upregulated (Fig. 6f-g). This dissociation demonstrates that *Rb* transcriptional quiescence is not a passive consequence of low abundance, but an actively maintained state enforced by the host vitellogenesis pathway. The upregulation of surface antigen genes *ompB* and *sca1*, which encode proteins involved in host-cell surface interactions in *Rickettsia* species (38), is consistent with symbiont niche reinforcement rather than pathogenic activation in this obligate endosymbiont. The host vitellogenesis pathway may represent one mechanism underlying the reduced conflict predicted between vertically transmitted endosymbionts and their hosts (39–41). The co-option of specific Vgr paralogs to implement symbiont quiescence during oocyte maturation exemplifies neofunctionalization (36) and suggests that the expanded Vgr repertoire represents an evolutionary investment in stable, low-conflict endosymbiont relationships (4, 20). How *Rb* suppresses host vitellogenesis genes remains unclear. This regulation may involve bacterial effector proteins, metabolic signaling, or indirect effects on host physiology. Identifying the bacterial factors responsible for this regulation will be an important next step.

While this study defines a multi-level regulatory framework connecting tick reproductive physiology to endosymbiont dynamics, several questions remain. First, although *Rb* DNA was detected in immature stages, we did not examine tissue localization or transcriptional activity in larvae or nymphs. Whether *Rb* also performs additional functions during immature development, including potential nutritional functions, remains to be determined. Second, the molecular basis of bidirectional regulation remains unclear. Whether this occurs through direct Vgr-symbiont interactions or indirect effects on the oocyte microenvironment cannot be resolved by RNAi alone and will require biochemical approaches. Third, only one Vgs and one Vgr candidate were functionally characterized in this study. Whether other members of the Vgs and Vgr families have similar regulatory functions remains to be determined. Finally, while basic genetic tools exist for Rickettsia species, precise gene editing in *Rb* is not currently feasible, limiting identification of specific bacterial factors involved in Vgs/Vgr suppression. Despite these limitations, the *I. scapularis-Rb* system provides a useful model for studying how blood-feeding arthropods regulate vertically transmitted endosymbionts. Much of what is known about reproductive symbiosis in arthropods comes from insect systems, such as *Wolbachia* (42). Our results show that ticks use a different biological approach, but may solve a similar problem: how to maintain a transmitted symbiont while limiting conflict with host reproduction. In this system, the vitellogenesis pathway appears to be one point where that balance is achieved.

## CONCLUSION

This study reveals a multi-layered regulatory system governing *Rb-I. scapularis* interactions: sex-dependent transcriptional silencing, ovary-specific localization, and vitellogenesis-mediated quiescence control. This study provides the first mechanistic explanation for endosymbiont regulation in blood-feeding ticks. We identified a critical dissociation between bacterial load and transcriptional state: Vgr12 controls symbiont abundance and transcriptional activity, while Vgs20 regulates transcriptional activity. This bidirectional regulation demonstrates that symbiont quiescence is actively enforced by the host, not a passive outcome of low bacterial numbers. The expansion of the Vgr gene family in *I. scapularis*, together with functional specialization of specific paralogs for endosymbiont regulation, represents a tick-specific strategy distinct from the largely single-copy Vgr systems found in other arthropods, including insects. This diversification likely reflects the unique reproductive and symbiotic biology of hard ticks, including obligate blood feeding and long-term maternal maintenance of obligate endosymbionts. These findings fill a critical gap in our understanding of vector-symbiont interactions and establish a foundation to study tick reproductive biology and vector competence in this medically important arthropod.

## MATERISALS AND METHODS

### Tick maintenance

*Ixodes scapularis* ticks were obtained from the Oklahoma State University Tick Rearing Facility and maintained as laboratory colonies at the University of Minnesota (UMN) and SUNY Upstate Medical University. At UMN, immature stages were blood-fed on hamsters (*Mesocricetus auratus*) and adults were fed using an artificial membrane feeding system with bovine blood (43–44). All animal procedures were approved by the University of Minnesota Institutional Animal Care and Use Committee (IACUC). Post-feeding females, unmated males, and mated males were separated for experiments. To generate *Rb*-depleted ticks, larvae were membrane-fed with ciprofloxacin (40 μg/ml)-containing blood and reared to adulthood. *Rb* depletion was verified by qPCR targeting the single-copy *gltA* gene (citrate synthase) in nymphs and the *Rb*-specific plasmid marker *ltrA2* (group I intron reverse transcriptase) in adults. Control ticks from the same cohort were fed antibiotic-free blood. Only verified *Rb*-positive and *Rb*-depleted adult females were used for subsequent experiments. At SUNY Upstate, colony ticks were used for molecular assays across all life stages, no blood-feeding procedures were conducted at this institution. Detailed protocols are provided in Supplementary Methods.

### Cell cultures and *R. buchneri* maintenance

ISE6 (*I. scapularis* embryonic) cells were cultured in complete medium (L-15C300 supplemented with 5% fetal bovine serum, 5% tryptose phosphate broth, and 0.1% lipoprotein concentrate) at 28 °C. *Rickettsia buchneri* ISO7^T^ (*Rb*) was maintained by serial passage in ISE6 cells in medium supplemented with 10% fetal bovine serum, 25 mM HEPES and 0.25% NaHCO₃, pH7 (14). To establish *Rb*-infected cultures, infected ISE6 cells were diluted 1:5 with uninfected cells and maintained at 28 °C. After 7-10 days, infection rate was determined by Giemsa staining (45) and cultures reaching ∼30% infected cells were used for experiments.

### Nucleic acid extraction and molecular analyses

Total DNA and RNA were extracted from tick cells, whole ticks across life stages, and dissected tissues using DNeasy Blood & Tissue Kit (Qiagen) and TRI Reagent (Sigma-Aldrich), respectively. RNA was purified by ethanol/isopropanol precipitation and cDNA synthesized using PrimeScript RT Reagent Kit with gDNA Eraser (Takara Bio). *Rb* detection across tick life stages was performed by PCR targeting *gltA* and *ltrA2* genes, with products resolved on 1.2% agarose gels. Gene expression was quantified by RT-qPCR (Bio-Rad CFX96) using SYBR Green detection. Seven *Rb* genes (*gltA, ltrA2, dnaK, rpoB, groEL, sca1, ompB*) were assessed, normalized to tick glyceraldehyde-3-phosphate dehydrogenase gene (*gapdh*) as housekeeping gene (45), and calculated using the 2^−ΔΔCt^ method. *Rb* genomic copy number was determined by quantitative PCR (qPCR) targeting the single-copy *gltA* gene using a standard curve. RNA/DNA ratios were calculated by quantifying both RNA copies (from cDNA) and DNA copies (from genomic DNA) for *gltA* using standard curves, expressed as RNA: DNA ratio per sample. Expression of *vgs20* (vitellogenin synthesis) and *vgr12* (vitellogenin receptor) genes was quantified across all tick life stages using RT-qPCR and normalized to tick *gapdh*. All primers are listed in Supplementary Table S1. All qPCR and RT-qPCR assays included no-template/no-RT controls, and specificity was verified by melt-curve analysis. Replication is reported in the figure legends.

### Transmission electron microscopy

Dissected tissues from unfed and engorged females were rinsed with PBS and processed for TEM analysis. Tissues were fixed in 2.5% glutaraldehyde, post-fixed in 2% OsO₄, dehydrated through graded ethanol series, and embedded in Embed 812 resin. Ultra-thin sections were stained with uranyl acetate and lead citrate and examined using a JEOL JEM-1400Plus transmission electron microscope. To quantify *Rb* abundance in developing oocytes, early-stage oocytes (oo I-II) were identified in sections from unfed and engorged females (n = 15 sections per group), and *Rb* cells were enumerated within size-matched oocytes. Detailed protocols are provided in Supplementary Methods.

### DNA Fluorescence In Situ Hybridization (FISH)

Tissue localization of *Rb* in dissected ovaries, salivary glands, and midguts from unfed females was examined by DNA FISH. An *ltrA2*-specific Cy3-conjugated probe unique to *Rb* among known *Rickettsia* species, was designed and validated in *Rb*-infected ISE6 cells before tissue application. Dissected tissues were fixed in Carnoy’s fixative, permeabilized, and hybridized overnight with the Cy3-labeled probe in hybridization buffer containing formamide. Following washes, nuclei were counterstained with DAPI and fluorescence visualized using a Leica SP8 confocal microscope. Probe sequence and detailed protocols are provided in Supplementary Methods.

### Genomic characterization and phylogenetic analysis of vitellogenesis genes

Putative vitellogenin synthesis (Vgs) and vitellogenin receptor (Vgr) genes in *I. scapularis* were identified by BLAST searches against the NCBI database using characterized *Drosophila melanogaster* sequences as queries. A total of 20 Vgs and 15 Vgr candidate genes were identified. Homologous sequences from representative tick species and arthropods were retrieved for comparative analysis. Protein sequences were aligned using MAFFT and maximum likelihood phylogenetic trees constructed using IQ-TREE with best-fit substitution models (46–48). Branch support was assessed using ultrafast bootstrap and SH-aLRT (49). Protein domain architectures were annotated using InterPro and Pfam databases, subcellular localization predicted using DeepLoc, and gene structures visualized using GSDS (50–52). For each aligned gene family, pairwise amino acid identity was calculated as the proportion of identical residues over aligned positions excluding gap-gap sites, generating 190 comparisons for Vgs (n = 20) and 105 for Vgr (n = 15). Mean pairwise identity was then calculated for each sequence by averaging its identity values against all other family members. Pairwise identity distributions between Vgs and Vgr families were compared using a two-sided Mann-Whitney U test. Detailed bioinformatics protocols are provided in Supplementary Methods.

### RNA interference (RNAi)

siRNA sequences targeting *Vgs20* (NCBI Reference sequence: XP_029841397.2) and *Vgr12* (NCBI Reference sequence: XP_042143231.1) were designed based on *I. scapularis* gene sequences and synthesized by Sigma-Aldrich. Scrambled siRNA (SsiRNA) served as negative control. For *in vitro* experiments, *Rb*-infected and uninfected ISE6 cells were transfected with 80 nM siRNA using Lipofectamine 3000 (Thermo Fisher Scientific, Waltham, MA, USA) (3 treatment groups: SsiRNA control, Vgs20 siRNA, Vgr12 siRNA; n=3 wells per group). For *in vivo* experiments, unfed adult females (n=18 per group for 3 treatment groups: SsiRNA control, Vgs20 siRNA, Vgr12 siRNA) were microinjected with 0.1 μM siRNA (100 nl volume/ per tick) into the opisthosoma.

Cells and ticks were harvested 5 days post-treatment for RNA extraction and qPCR analysis. Knockdown efficiency was confirmed by RT-qPCR normalized to *gapdh*. *Rb* genomic copy number and transcriptional responses were quantified as described above and expressed as log₂ fold change relative to controls. All siRNA sequences are listed in Supplementary Table S1. As a negative control, we used the MISSION® siRNA Universal Negative Control #1 (Sigma-Aldrich, Cat. No. SIC001), a non-targeting siRNA validated by the manufacturer to have no significant homology to any known human, mouse, or rat gene and to produce minimal effects on gene expression. The exact sequence is proprietary and not disclosed by the supplier.

### Statistical analysis

All statistical analyses were performed using GraphPad Prism (GraphPad Software, Boston, MA, USA). Group comparisons of detection rates used Fisher’s exact test. Multi-group gene expression comparisons used Kruskal-Wallis test with Dunn’s post hoc correction. TEM-based quantification of *Rb* abundance in oocytes was analyzed using a two-tailed Mann-Whitney U test. RNAi experiments were analyzed using a two-tailed Student’s t-test. Differences were considered significant at p < 0.05. Data are presented as mean ± SEM.

## CONFLICTS INTEREST

The authors declare that they have no conflict of interests.

## ACKNOWLEDGMENTS

We thank Gail Celio from the University Image Centers at the University of Minnesota, Twin Cities, for assistance with electron microscopy. Confocal images were acquired using instruments in the Upstate Medical University Neuroscience Microscopy Core facility. We particularly thank Scott Neal and Evelyn Voura from the Neuroscience Microscopy Core for their expert assistance with confocal microscopy. We also thank Dr. Yetrib Hathout from the School of Pharmacy and Pharmaceutical Sciences, Binghamton University, for valuable feedback and suggestions on the manuscript. The study was financially supported by a grant to X.R.W. from the SUNY Upstate Medical University start-up fund, Hendricks Pilot award, and a grant to U.G.M. from generous funding provided by grant R01AI049424 from the National Institutes of Health.

## Supplementary Figure legends

**Fig. S0. *Rb* transcription normalized to bacterial genome copy number.** *gltA* RNA/DNA ratios were determined in four DNA-positive male and four DNA-positive female ticks by RT-qPCR (RNA) and qPCR (DNA). Because *gltA* is a single-copy gene in the *Rb* genome, the RNA/DNA ratio provides an estimate of transcriptional activity per bacterium. The y-axis shows the log10-transformed RNA/DNA ratio. Each dot represents one biological replicate. Horizontal bars indicate the mean ± SEM. P < 0.01 (unpaired two-tailed Student’s t-test).

**Fig. S1. Ultrastructure of salivary glands in unfed and engorged female *I. scapularis*.** (**a-d**) TEM images of salivary gland acini from unfed female ticks showing acinar cell types I-III. Panels (**1–5**) show higher magnification views of selected regions illustrating typical salivary gland ultrastructure, including epithelial cells (EPC), secretory granules (sg), condensed secretory granules (csg), microvilli (mv), and intraductal lumen (ibd). No *Rb*-like bacteria were observed. (**e-f**) Salivary gland acini from fully engorged females showing late-stage glandular degeneration features in panels **6-8**: enlarged secretory compartments, reduced cellular organization, expanded secretory spaces, and numerous mitochondria (mi). No *Rb*-like bacteria were detected.

**Fig. S2. Ultrastructure of midgut tissues in unfed and engorged female *I. scapularis*.** (**a-b**) TEM images of midgut epithelial cells from unfed female ticks showing typical digestive cell morphology, including nuclei (n), mitochondria (mi), pigment granules (pg), and basal lamina (bl). No *Rb*-like bacteria were observed. (**c-e**) Midgut epithelial cells from engorged female ticks showing expanded digestive cells and organelles associated with blood digestion, including rough endoplasmic reticulum (rER), pigment granules (pg), and mitochondria (mi). No *Rb*-like bacteria were detected.

**Fig. S3. Cross-species phylogenetic analysis of Vgs genes across tick species.** ML phylogenetic tree of Vgs proteins from *I. scapularis* (Is) and related tick species. Collapsed clades are labeled with copy number (N) and species count (Spp). *Is* Vgs20 is highlighted. The inset table summarizes clade size, bootstrap support (SH-aLRT/UFBoot), and species composition for each clade. Species abbreviations: Is, *Ixodes scapularis*; Ir, *Ixodes ricinus*; Ipe, *Ixodes persulcatus*; Ip, *Ixodes pacificus*; Aa, *Amblyomma americanum*; Rm, *Rhipicephalus microplus*; Hl, *Haemaphysalis longicornis*. Tree scale bar represents substitutions per site.

**Fig. S4. Cross-species phylogenetic analysis of Vgr genes across tick species.** ML phylogenetic tree of Vgr proteins from *I. scapularis* and related tick species. Collapsed clades are labeled with copy number (N) and species count (Spp). Is Vgr12 is highlighted in the expanded Clade 9. The inset table summarizes clade size, bootstrap support, and species composition. Species abbreviations as in Fig. S3. Tree scale bar represents substitutions per site.

**Fig. S5. Comparative phylogenetic analysis of Vgs and Vgr families between *I. scapularis* and *D. melanogaster*.** (**a**) ML phylogenetic tree of Vgs proteins from *I. scapularis* (Is, red) and *D. melanogaster* (Dm, black). *I. scapularis* Vgs proteins form distinct, expanded clade relative to *Drosophila* homologs, indicating tick-specific gene family expansion. (**b**) ML phylogenetic tree of Vgr proteins showing pronounced divergence of tick Vgr repertoire relative to insect orthologs. Tree scale bars represent substitutions per site.

**Fig. S6. Broad comparative phylogenetic analysis of Vgs genes across arthropods, insects, and nematodes.** ML phylogenetic tree with collapsed clades color-coded by taxonomic group: green, *Ixodes*-restricted; orange, pan-insect; pink, Branchiopoda-restricted; tan, Diptera-restricted. Clades labeled with taxon pattern, copy number (N), and species count (Spp). Is Vgs20 highlighted. Inset table summarizes clade classification, copy pattern, size, species composition, bootstrap support, and taxonomic class. Species abbreviations used in this study: *Ixodes scapularis* (Is), *Anopheles gambiae* (Ag), *Apis mellifera* (Am), *Bombyx mori* (Bm), *Drosophila melanogaster* (Dm), *Daphnia pulex* (Dp), *Pediculus humanus* (Ph), and *Tribolium castaneum* (Tc).

**Fig. S7. Broad comparative phylogenetic analysis of Vgr genes across arthropods, insects, and nematodes.** ML phylogenetic tree with collapsed clades color-coded by taxonomic group and labeled with taxon pattern, copy number (N), and species count (Spp). Is Vgr12 highlighted within Clade 10. Inset table summarizes clade classification, copy pattern, size, species composition, bootstrap support, and taxonomic class. Species abbreviations: *Caenorhabditis elegans* (Ce); other abbreviations as in Fig. S6.

**Fig. S8. Gene structure analysis of Vgs and Vgr genes in *I. scapularis*.** (a) Exon-intron organization of all 20 Vgs genes drawn to scale (5’ to 3’). Red segments, exons; black lines, introns. Genomic span indicated on x-axis (kb). (b) Exon-intron organization of all 15 Vgr genes drawn to scale (5’ to 3’). Structure reveals variable gene architecture within families, with Vgr genes showing more complex intron-exon patterns than Vgs genes.

**Fig. S9. Protein domain architectures of Vgs and Vgr families in *I. scapularis*.** (**a**) Domain architectures of all 20 Vgs proteins showing pancreatic lipase-like domains as predominant feature. Protein lengths (aa) indicated at right. Domain annotations based on InterPro/Pfam database searches. (**b**) Domain architectures of all 15 Vgr proteins showing diverse domain compositions including LDLR-related, transmembrane, and other functional domains. Architecture diversity supports functional specialization within the Vgr family.

**Fig. S10. Conserved motif analysis of Vgs and Vgr proteins in *I. scapularis*.** (**a**) Conserved motif locations identified by MEME analysis across Vgs proteins. Colored blocks indicate positions of three conserved motifs; motif sequence logos and p-values shown at right. (**b**) Conserved motif locations across Vgr proteins.

